# CAT-APP: Contamination Analysis and Tempering—An Automated Online Platform for Plasma Proteomics with Data Rescuing Capabilities

**DOI:** 10.1101/2025.07.08.663798

**Authors:** Mingming Niu, Dong Zhang, Zhiwei Zhou, Jiaqi Zhang, Rong Zhang, Long Shen, Hong Wang

## Abstract

Plasma and serum proteome profiling have become central to biomarker discovery for medical research. The advent of high-throughput plasma proteomics pipelines has enabled the generation of massive datasets. However, biomarker discovery is frequently compromised by sample contamination, primarily from erythrocytes, platelets, and coagulation factors. Such contamination can lead to the false identification of contaminants as biomarkers in over 50% of studies or obscure genuine biomarkers due to contamination-induced variability. Currently, no tools are available to salvage such compromised data, and the default approach is often to discard the affected samples. To address this gap, we introduce CAT-APP (**C**ontamination **A**nalysis and **T**empering-An **A**utomated Online Platform for **P**lasma **P**roteomics). CAT-APP tackles contamination through three key modules: multi-dimensional contamination assessment and adaptive contamination indexing, mathematical model-based contamination tempering, and data recovery evaluation with visualization. We applied CAT-APP to 28 independent plasma proteomics datasets and found that 82% exhibited contamination issues. CAT-APP markedly improved data quality by removing false-positive biomarkers introduced by contamination and restoring true biological signals that were previously obscured in representative datasets. To promote widespread adoption, we provide CAT-APP as an freely accessible, user-friendly, and robust web platform at https://www.bloodecosystem.com/tools/CAT-APP/.

## Introduction

Plasma and serum, collectively referred to as plasma for simplicity, are readily accessible bodily fluid that contains molecular signatures reflective of both physiological and pathological states, making them the most widely used specimen in clinical diagnostics ^1,2^. The discovery of protein biomarkers through plasma proteome profiling has been a long-standing goal in biomedical research ^3–5^. Recent advances in high-throughput technologies have led to the generation of large volumes of plasma proteomic data ^6,7^. These include mass spectrometry-based platforms, aptamer-based assays (e.g., SomaLogic), and antibody-mediated assays (e.g., Proximity Extension Assay, PEA) ^8–10^. For example, the UK Biobank has invested over billions of dollars in large-scale plasma proteome profiling of its cohort ^11^. Despite these technological strides and significant financial investment, clinical practice remains limited by a scarcity of validated biomarkers, most of which were discovered decades ago ^12,13^.

The discovery of plasma biomarkers faces numerous challenges, with sample contamination being among the most significant. This contamination primarily arises from three sources: erythrocytes, due to hemolysis; platelets, resulting from inconsistencies in centrifugation speed and duration; and coagulation factors, influenced by variations in anticoagulant types and coagulation times ^12,14,15^. These sources of contamination often stem from inconsistencies in sample collection, processing, and storage procedures, especially in multi-center cohort studies ^16^. In clinical settings, where patient care typically takes precedence over strict adherence to sample protocols, it is particularly difficult to fully eliminate such contamination ^17^. Consequently, contaminant proteins may be falsely identified as biomarkers, or genuine biomarkers may be obscured due to increased variability. Notably, over half of plasma proteomic biomarker studies have identified biomarker candidates that are contaminant proteins ^14^.

Previous studies have proposed panels of biomarkers to assess contamination levels in plasma samples, enabling the exclusion of compromised samples to reduce the risk of falsely identifying contaminant proteins as genuine biomarkers ^14^. However, contamination is pervasive in samples collected from hospitals and biobanks. While this exclusion-based approach can help mitigate false discoveries, it cannot recover true signals obscured by contamination-induced variability. More importantly, discarding affected samples results in the loss of valuable biospecimens and incurs substantial costs due to the proteomic profiling of samples that are ultimately unusable ^18,19^. Currently, no tools are available to correct these confounding effects and salvage compromised datasets. Consequently, many studies either remain unaware of the severity of this issue or continue to discard contaminated samples by default.

To bridge this gap, we introduce **CAT-APP**: an automated online platform for plasma proteomics with built-in data rescuing capabilities. CAT-APP integrates three key modules: multi-dimensional contamination assessment and adaptive contamination indexing, mathematical model-based contamination tempering, and data recovery evaluation with visualization. Using CAT-APP, we analyzed 28 independent plasma proteomics datasets and found that 82% exhibited contamination issues. We further demonstrate CAT-APP’s effectiveness in rescuing data by eliminating false-positive biomarkers caused by contamination and restoring true biological signals that were previously obscured. CAT-APP is provided as a freely accessible, user-friendly, and robust online platform, available at https://www.bloodecosystem.com/tools/CAT-APP/, to facilitate broad adoption.

## Results

### Blood contamination severely confounds plasma proteome studies

In a recent large-cohort plasma proteomic study investigating acute infections, we unexpectedly identified a substantial subset of outlier samples characterized by markedly elevated expression of a series of proteins. These aberrant expression patterns significantly confounded the overall cohort data, rendering these samples clear outliers (Fig. 1A). Notably, the distinction between these outlier samples and the rest of the cohort was even more pronounced than the differences observed between infected individuals and healthy donors (HD) (Fig. 1B). To investigate the source of this data distortion, we analyzed the overly expressed proteins in the outlier samples. These proteins exhibited exceptionally high inter-protein correlations **(Fig. 1C, Supplementary Fig. S1A)** and elevated CV compared to other proteins (Fig. 1D), suggesting they are co-affected by a shared confounding factor that differs markedly between outlier and non-outlier samples. Pathway analysis revealed strong enrichment in platelet-related pathways, rather than disease-associated pathways (Fig. 1E). Notably, 98.7% of these suspect proteins overlapped with the top 50% of the most abundant proteins in published platelet proteome (Fig. 1F, Supplementary Fig. S1B), indicating insufficient removal of platelets from these plasma samples. Cross-referencing with published platelet contamination markers revealed a strong overlap with the top proteins enriched in the outlier group (Fig. 1G). Principal component analysis (PCA) using experimentally validated platelet markers effectively separates all outlier samples, mirroring the separation observed with the full plasma proteome (Supplementary Fig. S1C). Together, these findings support the conclusion that the observed distortions are primarily attributable to platelet contamination ^14^. Retrospective analysis further traced the source of contamination to changes in centrifugation speed and duration, which had been modified by clinical staff on a specific collection day to accommodate an unexpectedly high sample volume (Fig. 1H).

**Figure 1.**
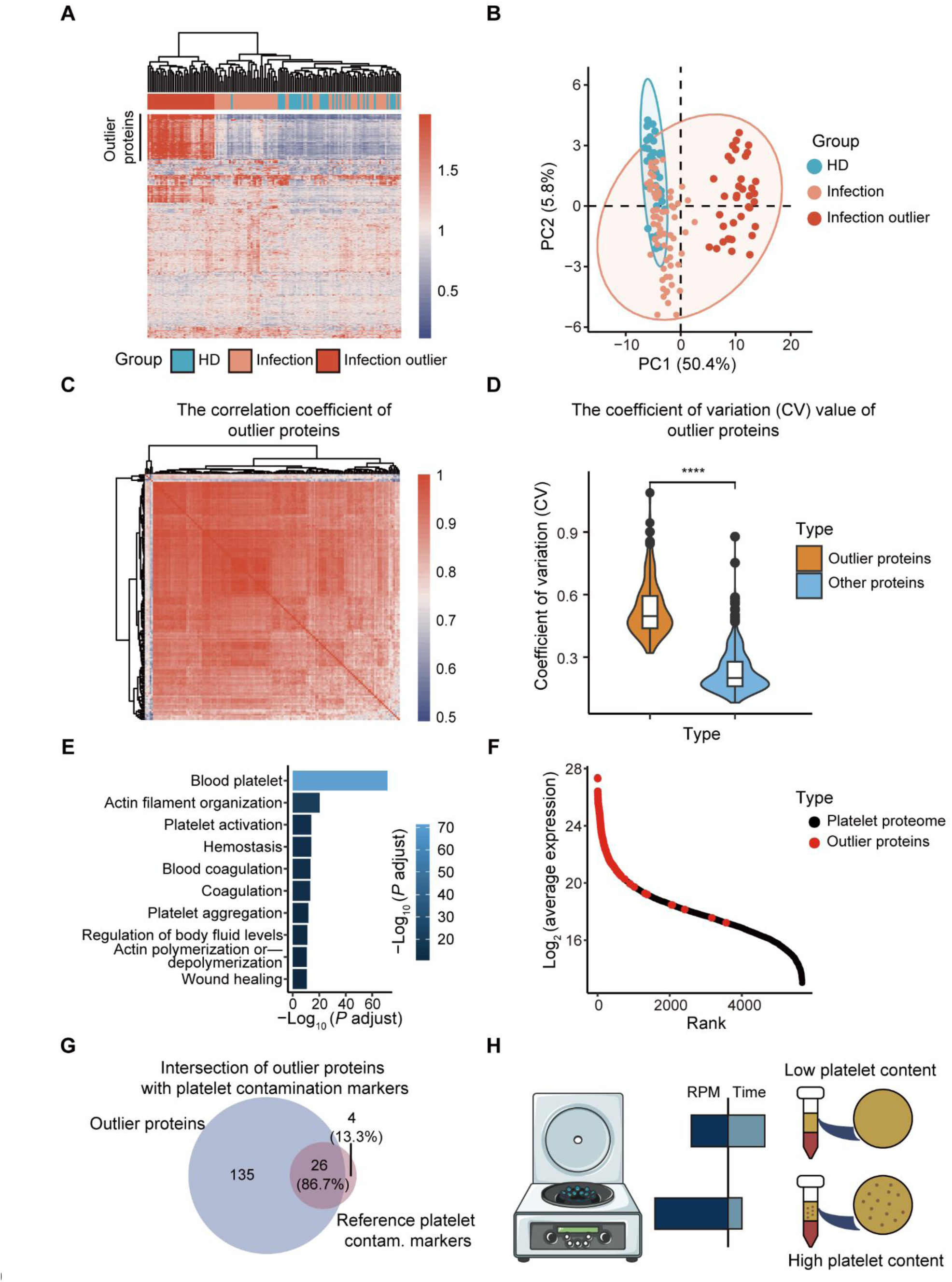
Platelet contamination severely confounds plasma proteome study. (**A**) Heatmap of plasma proteomic profiles showing a distinct subset of outlier samples with aberrantly elevated protein expression. (**B**) PCA reveals clear segregation of outlier samples. (**C**) Pairwise correlation matrix illustrates strong inter-protein correlation within the outlier proteins. (**D**) Violin plot comparing CV between outlier proteins and all other proteins (P value = 2.22e-16, Wilcoxon rank-sum test). (**E**) Gene Ontology enrichment analysis of the aberrantly expressed proteins reveals significant enrichment of platelet-related pathways (top 10 terms by adjusted P value). (**F**) Rank plot showing the rank of outlier proteins (red) in published platelet proteome. (**G**) Venn diagram comparing outlier proteins with experimentally validated platelet contamination (contam.) protein markers. (**H**) Schematic illustration showing that platelet contamination is attributable to variations in centrifugation speed and duration during blood sample processing.

### Contamination maker-based correction model recovers affected samples

In clinical settings, patient care often takes priority over strict sample handling, making contamination hard to avoid and common. While removing affected samples can reduce false discoveries, it also wastes valuable clinical resources, diminishes statistic power, and increases costs due to discarded profiling efforts. Thus, to address the contamination issue and preserve valuable data, we instead developed a workflow incorporating a mathematical model to recover affected data rather than discard it (Fig. 2A).

**Figure 2.**
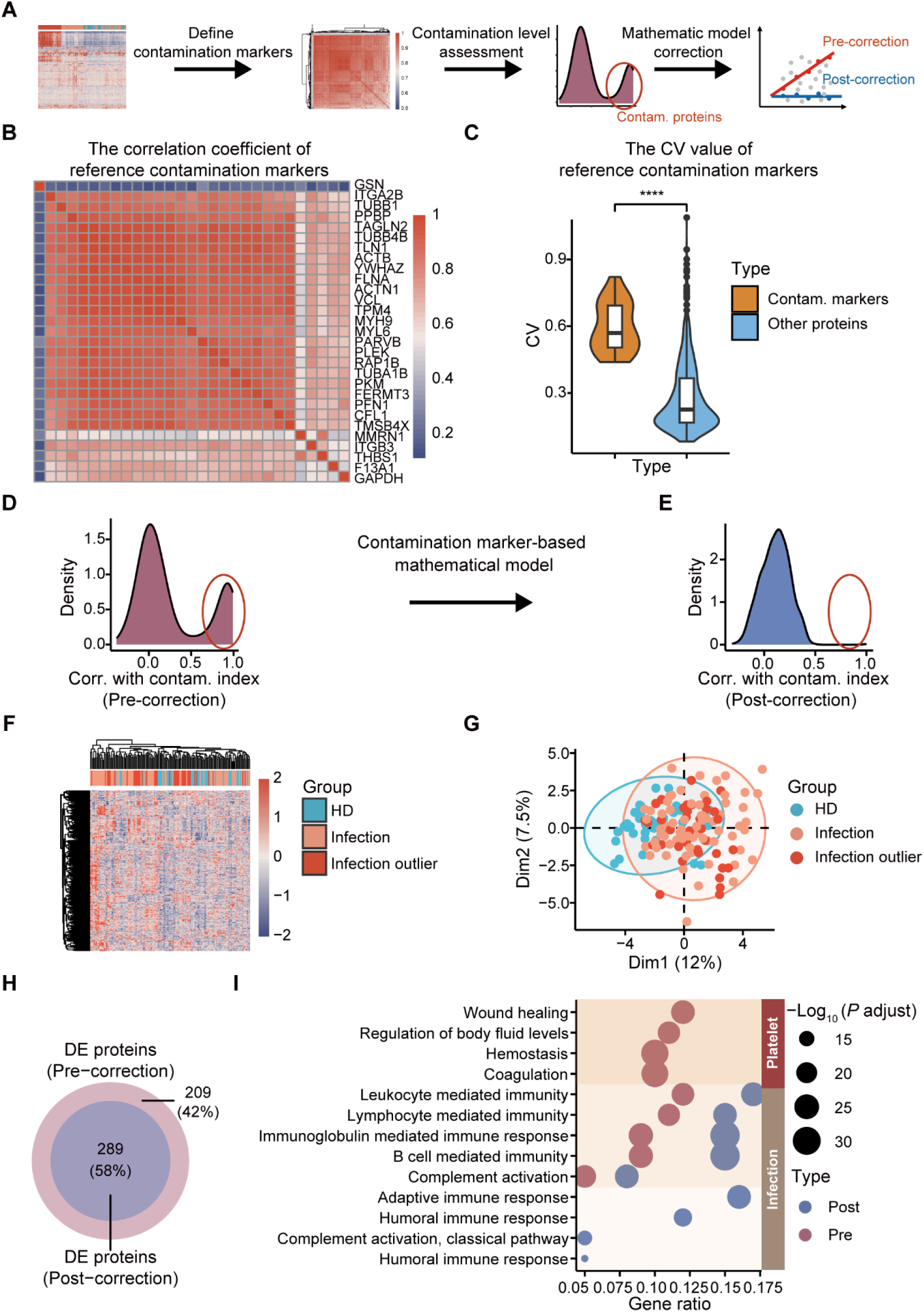
Contamination marker-based correction model restores data quality in affected samples. (**A**) Schematic overview of the contamination correction workflow. (**B**) Pairwise Pearson correlation coefficients among reference contamination protein markers ^14^. (**C**) Violin plot comparing CV distributions between contamination markers and other plasma proteins (P < 5.5e-13, Wilcoxon rank-sum test). (**D**) Density plot showing the distribution of correlations between plasma proteomes and the contamination index before correction (**E**) Density plot showing the distribution of correlations between plasma proteomes and the contamination index after correction. (**F**) Heatmap of corrected proteomic data illustrating restoration of outlier samples. (**G**) PCA plot showing recovery of outlier samples following correction. (**H**) Venn diagram of DE proteins identified before and after correction. (**I**) Comparison of pathway enrichment results (top 10 terms) before and after correction.

We first applied an experimentally validated panel of platelet contamination markers as a reference to assess contamination levels across the cohort samples ^14^. We hypothesized that if platelet contamination varies substantially among samples, and if the expression of these marker proteins is primarily driven by contamination rather than true biological differences, then these marker proteins should exhibit strong inter-correlations and high variability across samples. As expected, we observed both strong inter-correlations among the contamination markers and significant variability in their expression levels among samples (Fig. 2B and C).

Recognizing that the expression of certain markers may be influenced by both contamination and genuine biological changes depending on the context, we refined our selection by including only those platelet biomarkers with high inter-correlation (r > 0.9) as study-specific contamination indicators. To further identify proteins affected by contamination, we calculated the mean contamination level of the selected contamination markers, defining this as the contamination index. We then performed a correlation analysis between the contamination index and all detected plasma proteins. Theoretically, in the absence of contamination, the distribution of protein correlations with contamination index should follow a Gaussian distribution centered around zero. However, we observed a clear bimodal distribution, with a substantial subset of proteins showing strong positive correlations with the contamination markers, indicating widespread contamination effects (Fig. 2D).

Next, we implemented a mathematical model–based strategy to correct protein-level contamination using each protein’s correlation with the contamination index. This approach effectively removed the influence of platelet contamination (Fig. 2E, Supplementary Fig. S1D). Following correction, the outlier effects in contaminated samples were eliminated, revealing a trend of separation between disease groups and healthy donors that were previously masked by platelet contamination (Fig. 2F and G). Differential expression (DE) analysis between disease groups and healthy donors identified 498 DE proteins before correction. After applying the correction, 209 DE proteins were removed, including most of the outlier contamination proteins (Fig 2H, Supplementary Fig. S1E and S1F). Pathway analysis comparing pre- and post-correction results revealed that platelet contamination-related pathways were successfully eliminated, while enrichment of infection-related pathways, such as those involved in inflammatory and immune responses, became more prominent. Importantly, several biologically meaningful changes that were previously masked by platelet contamination, such as those related to the adaptive immune response, were also uncovered (Fig. 2I). Together, these results demonstrate that the dynamic correction model we developed can effectively recover valuable plasma proteomic data affected by platelet contamination, avoiding the need to discard compromised samples.

### CAT-APP: an accessible, user-friendly, and robust online platform for plasma proteome studies

Currently, no dedicated tools exist to address contamination in plasma proteomics data. To fill this gap, we developed CAT-APP (**C**ontamination **A**nalysis and **T**empering-An **A**utomated Online Platform for **P**lasma **P**roteomics), a robust, user-friendly, and freely accessible web platform designed to support wide adoption in the proteomics community (Fig. 3).

**Figure 3.**
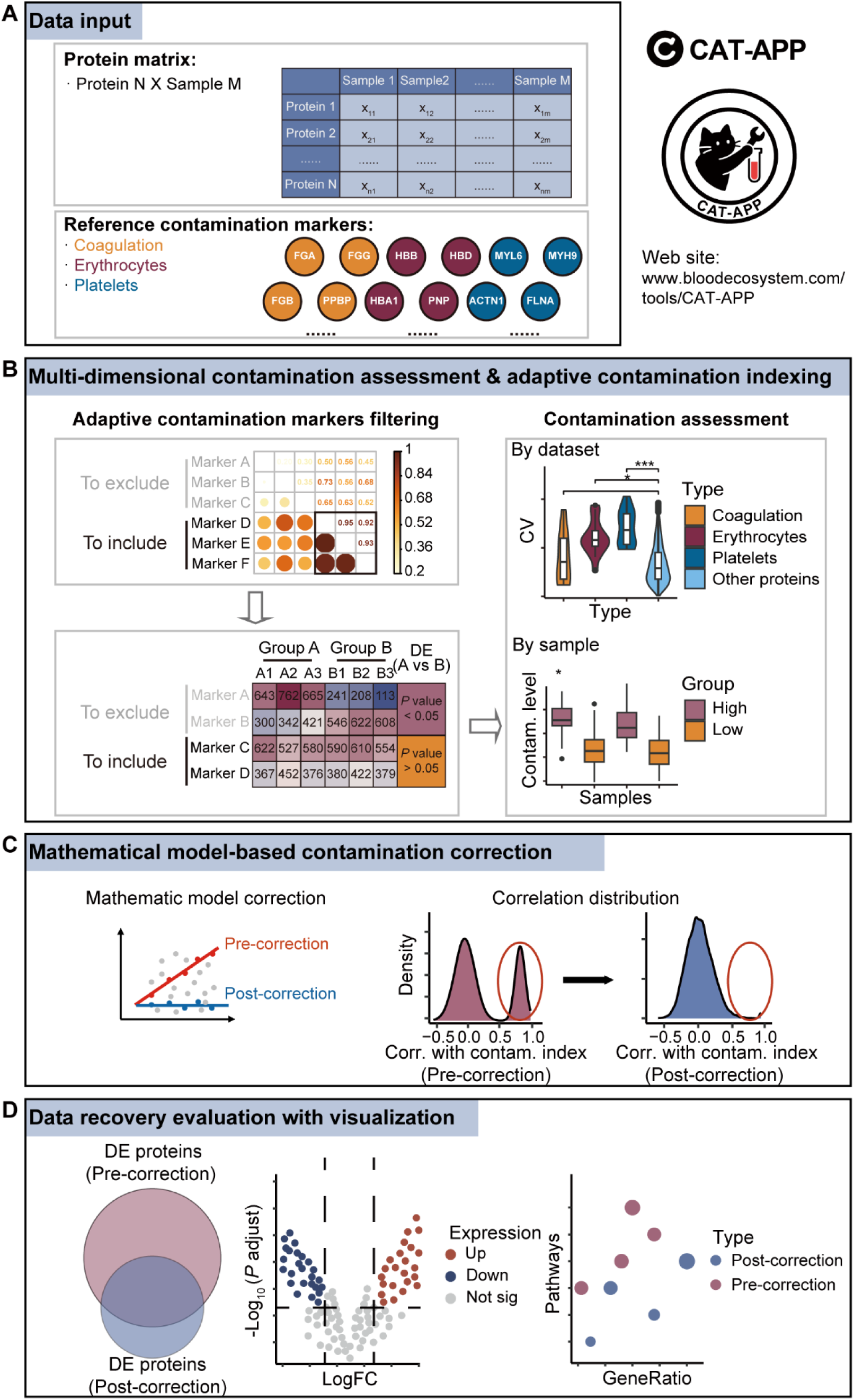
CAT-APP is designed as an accessible online platform for plasma proteomics analyses. (**A**) Illustration of the data input module. Users can upload a protein abundance matrix along with optional sample group annotations. The platform includes experimentally validated contamination marker panels for erythrocytes, coagulation, and platelets, with support for user-defined extensions. (**B**) Multi-dimensional contamination assessment and adaptive contamination indexing module. Users can select study-specific contamination markers based on high correlation, with an option to exclude markers that show significant differences between sample groups. The overall contamination level is further evaluated using the difference in CV between contamination markers and the rest of the proteome. (**C**) Contamination correction module based on a mathematical model. Visualizations of the data distributions before and after correction relative to the contamination index. (**D**) Data recovery evaluation and visualization module. Supports DE analysis with visual outputs of a volcano plot comparing pre- and post-correction DE proteins. By integrating pathway enrichment analyses, data recovery evaluation can be further visualized through pathway enrichment scatter plots.

The platform integrates three core modules: (1) multi-dimensional contamination assessment and adaptive contamination indexing, (2) mathematical model-based contamination correction, and (3) data recovery evaluation with visualization. CAT-APP accepts a protein expression matrix as input, ensuring compatibility with diverse proteomics pipelines. Users can upload their data and optionally assign sample groupings for downstream dynamic contamination marker selection and differential expression analysis. To systematically address major sources of plasma contamination, CAT-APP includes experimentally validated reference markers for platelets, erythrocyte hemolysis, and coagulation ^14^ (Fig. 3A).

CAT-APP employs an adaptive, rather than fixed, set of contamination markers because the expression of certain markers may reflect both contamination and true biological variation, depending on the study context. To identify study-specific reliable markers for correction, CAT-APP first evaluates pairwise correlations among reference contamination markers. This approach is based on the assumption that if contamination is the primary source of variation, most markers will exhibit high pairwise correlation. Conversely, low correlation suggests minimal contamination influence. By default, markers with pairwise correlation coefficients greater than 0.9 are selected for correction; Users may adjust this correlation threshold to fit the goals of their study (Fig. 3B).

Secondly, contamination marker-based correction model can inadvertently remove true biological signals if contamination levels are not randomly distributed but are instead systematically biased across sample groups. For example, if contamination marker levels are consistently higher in disease samples and lower in healthy controls, the model may misattribute genuine biological differences to contamination, leading to the unintended removal of true signals. To mitigate this, CAT-APP performs differential expression analysis of contamination markers between user-defined biological groups. Markers that are significantly different between groups are excluded from the following correction process. CAT-APP also evaluates whether contamination markers show greater variability than the background plasma proteome to estimate the extent of contamination and determine whether correction is warranted (Fig. 3B). Additionally, to enhance correction accuracy and avoid model degradation due to weakly correlated markers, CAT-APP restricts correction to the top 30% of contamination markers most strongly correlated with each protein, using a minimum of three markers.

After adaptive contamination indexing and completing contamination assessment, CAT-APP calculates the mean contamination of the selected markers to derive a final contamination index. Using this index, CAT-APP performs mathematical model-based contamination correction (see Materials and Methods) to adjust for contamination and recover affected protein signals. The platform provides visualizations that compare the distribution of correction effects across all plasma proteins in relation to the contamination index, enabling users to assess the outcome. A successful correction is typically reflected by the transformation of a right-skewed or bimodal distribution into a near-normal distribution (Fig. 3C).

Finally, CAT-APP enables differential expression analysis between user-defined biological groups following contamination correction. Combined with downstream pathway analyses, these features help evaluate CAT-APP’s effectiveness in removing false-positive biomarkers caused by contamination and recovering true biological signals previously obscured (Fig. 3D). CAT-APP is freely accessible at: https://www.bloodecosystem.com/tools/CAT-APP/.

### CAT-APP reveals widespread contamination in plasma proteome studies and rescues affected datasets

To systematically evaluate CAT-APP’s performance, we applied the platform to several published plasma proteome datasets. In a neurodegenerative disease cohort, CAT-APP identified erythrocyte-derived contamination, as evidenced by substantially higher CV in erythrocyte-associated markers compared to the global plasma proteome, along with right-skewed correlation distributions between protein abundance and the erythrocyte contamination index (Fig. 4A and B). After CAT-APP correction, the correlation distribution approximated normality, and the elevated CVs of erythrocyte markers were markedly reduced (Fig. 4C and D). DE analysis between the neurodegenerative disease and healthy control groups revealed an increase in DE proteins from 401 (pre-correction) to 605 (post-correction), representing a ∼50% gain. Pathway analysis further confirmed the removal of erythrocyte-related signatures, such as those associated with reductase activity, following correction ^20^. Notably, biologically relevant pathways that had been previously masked by contamination, such as those linked to neurodegeneration and Alzheimer’s disease, became significantly enriched post-correction, underscoring CAT-APP’s ability to recover meaningful biological signals (Fig. 4E and F).

**Figure 4:**
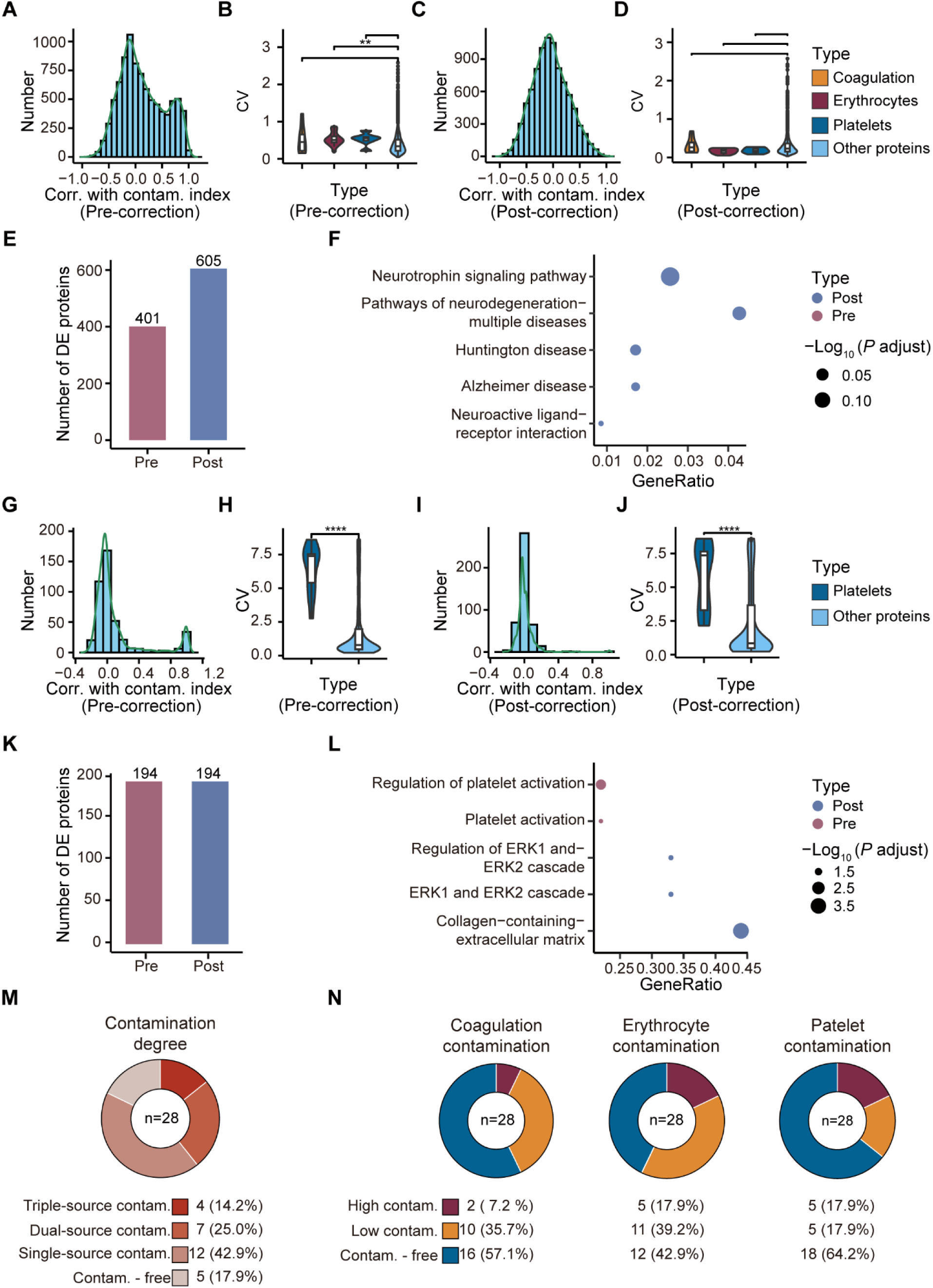
CAT-APP reveals widespread contamination in external plasma proteome datasets and rescues affected datasets. (**A**) Density plot showing the distribution of correlations between the plasma proteome and the contamination index in a representative neurodegenerative disease study, prior to correction. (**B**) Comparison of CV between contamination marker panels (coagulation, erythrocyte, and platelet) and the rest of the plasma proteome prior to correction. Erythrocyte markers show a significantly higher CV (P = 0.004, Wilcoxon rank-sum test). (**C**) Density plot showing the distribution of correlations between the plasma proteome and the contamination index after correction. (**D**) Comparison of CV between contamination marker panels (coagulation, erythrocyte, and platelet) and the rest of the plasma proteome after correction. The previously elevated CV of erythrocyte markers is effectively reduced following correction. (**E**) Bar plot illustrating the number of DE proteins pre- and post-correction. (**F**) Comparison of pathway enrichment results using DE proteins before and after correction reveals newly identified pathways that are highly functionally relevant to neurodegenerative disease. (**G**) Density plot showing the distribution of correlations between the plasma proteome and the contamination index in a representative cardiomyopathy disease study, prior to correction. (**H**) Comparison of CV between platelet contamination marker panels and the rest of the plasma proteome prior to correction. Platelet markers show a significantly higher CV (P value = 3.1e-5, Wilcoxon rank-sum test). (**I**) Density plot showing the distribution of correlations between the plasma proteome and the contamination index after correction. (**J**) Comparison of CV between platelet contamination marker panels and the rest of the plasma proteome after correction. (**K**) Bar plot illustrating the number of DE proteins pre- and post-correction. (**L**) Comparison of pathway enrichment results using DE proteins before and after correction. Platelet contamination–associated pathways are successfully removed, while newly identified pathways are functionally related to cardiomyopathy. (**M**) Pie chart summarizing contamination detection across 28 datasets: only 5 datasets (17.9%) show no detectable contamination from any of the three sources. (**N**) Overview of contamination prevalence across 28 public datasets: 57.1% exhibit erythrocyte contamination, 42.9% coagulation, and 35.8% platelet-related contamination.

In a cardiomyopathy cohort with only moderate contamination, although the variability of platelet contamination markers was only partially reduced, and the number of differentially expressed proteins changed minimally after correction (Fig. 4G-K). Pathway analysis revealed successful removal of platelet-related signatures and enhanced enrichment of cardiomyopathy-relevant biological processes, such as the collagen-containing extracellular matrix ^14^ (Fig. 4L). These findings demonstrate CAT-APP’s ability to reduce contamination-driven bias and uncover meaningful biological signals, even in datasets without overt contamination artifacts.

To assess the prevalence of contamination in plasma proteomics, we systematically analyzed 28 published datasets from iProX, PRIDE, and PubMed. Contamination was identified in 82.1% of the datasets, with only 17.9% classified as contamination-free (Fig. 4M), underscoring the widespread nature of this issue in plasma proteomic studies. Among contaminated datasets, 42.9% exhibited single-source contamination (erythrocyte-, platelet-, or coagulation-related), while 39.2% showed evidence of multi-source contamination. To evaluate contamination severity, we compared the expression-level variability of contamination markers with that of non-marker proteins (Fig. 4N). Based on this analysis, datasets were stratified into three tiers: high contamination, low contamination, and contamination-free (See methods). Erythrocyte- and platelet-derived contamination were most prevalent among high-severity cases, each affecting 17.9% of datasets, whereas coagulation-related contamination was less common (7.2%). This hierarchy highlights the critical need for stringent standardization of plasma collection and preparation protocols. In addition, we observed that contamination markers do not exhibit consistently high correlations across contaminated datasets. This variability reflects the complexity of contamination sources and the case-specific characteristics of individual studies, underscoring the importance of the adaptive contamination indexing function that enables CAT-APP to perform optimally across diverse datasets (Supplementary Fig. S2). Given the practical challenges of fully eliminating contamination in routine clinical settings, the pervasive contamination observed across repositories (82.1%) emphasizes the urgency for robust correction tools and highlights the broad utility of CAT-APP in addressing this critical gap in plasma proteomics research.

## Discussion

Plasma proteomics holds tremendous promise for biomarker discovery in clinical diagnostics and biomedical research due to plasma’s accessibility and its rich molecular content reflective of physiological and pathological states. However, widespread contamination from erythrocytes, platelets, and coagulation factors has emerged as a critical bottleneck in the field. Critical factors include centrifugation speed, anticoagulant type ^21^, blood collection tube selection ^22^, which may trigger platelet activation ^23^ or complement system activation ^24^. Additionally, sample hemolysis has been shown to adversely affect mass spectrometry analysis ^25,26^, and variable biobank storage conditions can introduce further bias ^27^. Our systematic analysis revealed that over 80% of published plasma proteomics datasets harbor significant contamination, underscoring the pervasive nature of this problem. Contamination not only leads to false-positive biomarker identification but also obscures genuine biological signals, thereby hindering clinical translation and discovery efforts.

Most previous plasma proteomics studies did not fully recognize the severity of contamination. Even when acknowledged, the standard practice has typically relied on excluding contaminated samples using predefined marker panels. Although this approach reduces false discoveries, it comes at the cost of losing valuable data and diminishing statistical power. Moreover, no publicly available tools currently exist to correct contamination effects and recover compromised datasets, leaving a critical gap in plasma proteomics workflows. CAT-APP addresses this unmet need by providing an integrated, automated, and user-friendly online platform that enables contamination detection, adaptive indexing, correction, and data recovery evaluation. This solution may represent a paradigm shift from exclusion-based to recovery-based strategies, allowing for more effective and informative utilization of plasma proteomics data.

The adaptive contamination indexing module of CAT-APP tailors the contamination signature to the dataset’s context, improving the accuracy of contamination quantification. This adaptability is crucial given the heterogeneity in sample collection, handling, and disease context observed across cohorts ^28^. Moreover, by excluding contamination markers that differ significantly between biological groups, CAT-APP mitigates the risk of inadvertently removing true biological signals during correction.

The mathematical correction model effectively removes contaminant effects while preserving genuine biological variation, as demonstrated in multiple representative cohorts. In an acute infection cohort, CAT-APP successfully identified and removed platelet contamination, unmasking biologically relevant immune response signatures that were previously obscured. Similarly, in neurodegenerative and cardiomyopathy cohorts, the platform reduced erythrocyte- and platelet-derived contamination signals, respectively, and enhanced the detection of disease-associated pathways. Notably, correction resulted in a substantial increase in the number of differentially expressed proteins, reflecting improved sensitivity and specificity of biomarker discovery post-correction.

Our survey across 28 external datasets further confirmed the widespread and heterogeneous nature of plasma contamination, with frequent multi-source contamination complicating data interpretation. This observation reinforces the critical need for standardized plasma handling protocols but also highlights the practical limitations of achieving contamination-free samples in routine clinical settings. CAT-APP’s capability to adaptively index and correct contamination across diverse datasets offers a practical solution to this challenge, maximizing the utility of existing plasma proteomic data and enhancing reproducibility across studies.

The availability of CAT-APP as a free, web-based platform lowers the technical barrier for adoption, enabling researchers from diverse backgrounds to implement robust contamination assessment and correction without requiring specialized computational expertise. The platform’s visualizations provide intuitive diagnostics of contamination levels and correction outcomes, facilitating transparent and informed decision-making. Moreover, CAT-APP’s modular design allows for future expansion to accommodate additional contamination sources and correction strategies as the field of plasma proteomics continues to evolve. Importantly, the CAT-APP framework may also be extended to other biofluids or even tissue samples where variability due to blood contamination is a concern, broadening its applicability beyond plasma and serum.

In conclusion, CAT-APP fills a critical void in plasma proteomics workflow by providing an accessible, adaptable, and effective tool for contamination management and data recovery. By enabling the rescue of compromised samples and enhancing the robustness of biomarker discovery, CAT-APP has the potential to accelerate clinical translation and advance the development of personalized medicine.

## Methods

### Data collection and preprocessing

A total of 30 plasma proteomics datasets were analyzed in this study, including 2 datasets previously reported in our publication and 28 publicly available datasets retrieved from iProX, and PRIDE. These datasets span a variety of clinical conditions, including neurodegenerative, infectious, and cardiovascular diseases. All datasets were preprocessed by imputing missing values using the seqknn method from the multiUS package, followed by log2 transformation. The resulting normalized data were then used for all downstream analyses.

### Multi-dimensional contamination assessment and adaptive contamination indexing

To identify dataset-specific contamination markers, we employed a dynamic selection strategy. Experimentally validated reference markers were compiled from a previous study ^14^. These markers were first filtered using pairwise Pearson correlation analysis, retaining only those with a mean correlation coefficient greater than 0.9 across the set. When sample group annotations were available, markers showing significant between-group differences (Wilcoxon rank-sum test, P < 0.05) were excluded to avoid conflating biological variation with technical artifacts. To further ensure marker robustness, we restricted the final marker panel to the top 30% of markers most strongly correlated with each other, while requiring a minimum of three markers, to prevent model degradation from weakly correlated features. The resulting high-confidence marker set was then used to calculate the contamination index, defined as the mean contamination level across the final panel. Contamination levels were classified into three categories: (i) Uncontaminated, defined as datasets lacking highly correlated contamination markers (Pearson correlation < 0.9); (ii) Low contamination, where highly correlated markers were present, but the coefficient of variation (CV) of contamination marker panel was not significantly higher than that of other proteins (P > 0.05); and (iii) High contamination, where both highly correlated contamination markers (Pearson correlation < 0.9) and significantly elevated marker CVs were observed (P < 0.05).

### CAT-APP web platform

The CAT-APP platform was developed using R (v4.4.2) and Shiny (v1.9.1). Users can upload a protein abundance matrix along with an optional group annotation file containing sample IDs and group labels. The platform performs automated quality control by evaluating pairwise correlations and CV distributions for platelet-, erythrocyte-, and coagulation-related markers. When group information is provided, CAT-APP additionally assesses between-group differences to avoid misattributing true biological variation to contamination. Based on the quality control results, the platform automatically reports whether contamination is present and whether data correction is recommended. Users can also customize correlation thresholds to refine quality control criteria and re-run the analysis as needed.

Following contamination correction, CAT-APP provides built-in modules for downstream differential expression and functional enrichment analysis. Users can perform group comparisons on the corrected protein matrix and visualize results through interactive plots, facilitating immediate biological interpretation of corrected data.

### Bioinformatics analysis

All statistical and visualization analyses were conducted in R (v4.4.2). Data manipulation employed tidyr and stringr. Visualization was accomplished using pheatmap, ggplot2, ggpubr, ggrepel, and VennDiagram. Differential expression analysis was performed with limma ^29^, and functional enrichment analysis utilized clusterProfiler ^30–32^. Correlation analyses used the Rfast and stats packages. Mathematical modeling was conducted via the MASS package.

## Acknowledgements

We thank the Center for Advanced Technologies at the Haihe Laboratory of Cell Ecosystem for their technical support.

This study was supported by grants from the CAMS Innovation Fund for Medical Science (2023-I2M-3-014 to H.W., 2023-I2M-2-007 to H.W., 2021-I2M-1-073 to H.W.), the National Natural Science Foundation of China (82341080 to H.W., 82270236 to H.W., 82400271 to M.N., 82273217 to L.S.), the Fundamental Research Funds for the Central Universities, Peking Union Medical College (3332024200 to M.N.), the distinguished Young Scholars of Tianjin (22JCJQJC00090 to H.W.), the China Foundation For Youth Entrepreneurship and Employment-Incaier Public Welfare Fund (HH25KYHX0004 to D.Z.), the Tianjin Municipal Science and Technology Commission Grant (22JCQNJC00040 to L.S.), and the Natural Science Foundation of Sichuan Province (2025ZNSFSC0737 to H. W.).

## Author contributions

H.W. and M.N. conceived the concept and supervised the overall study. H.W., M.N., and D.Z. designed the CAT-APP platform. D.Z., J.Z., and Z.Z. performed the data analysis with input from all authors. L.S. and R.Z. participated in study coordination and contributed to manuscript preparation. H.W., M.N., and D.Z. wrote the manuscript with input from all authors.

---

**Figure S1:**
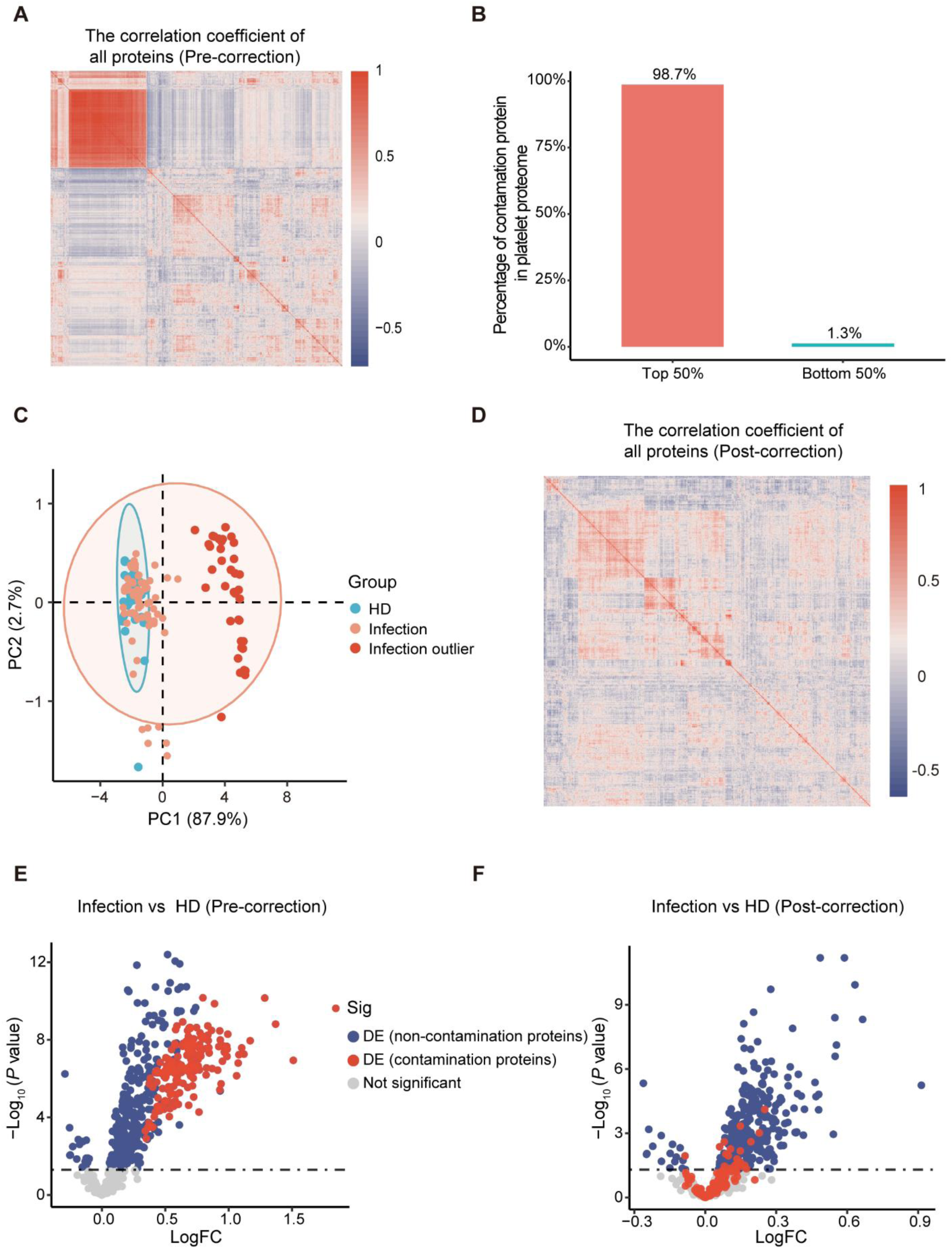
Characteristics of contamination and evaluation of correction using experimentally validated platelet contamination markers. (A) Pairwise Pearson correlation coefficients of all proteins. Contaminant proteins exhibit significantly stronger inter-protein correlations compared to other proteins. (B) 98.7% of contaminant proteins rank within the top 50% of the most abundant proteins in the platelet proteome. (C) PCA using experimentally validated platelet contamination markers highlights separation of outlier samples prior to correction. (D) Pairwise Pearson correlation coefficients of all proteins. The previously elevated inter-protein correlations among contaminant proteins are reduced after correction. (E) Volcano plot shows that all contaminant proteins are falsely identified as DE proteins prior to correction. (F) Post-correction volcano plot shows that most of contaminant proteins, previously misidentified as DE, are no longer classified as DE.

**Figure S2:**
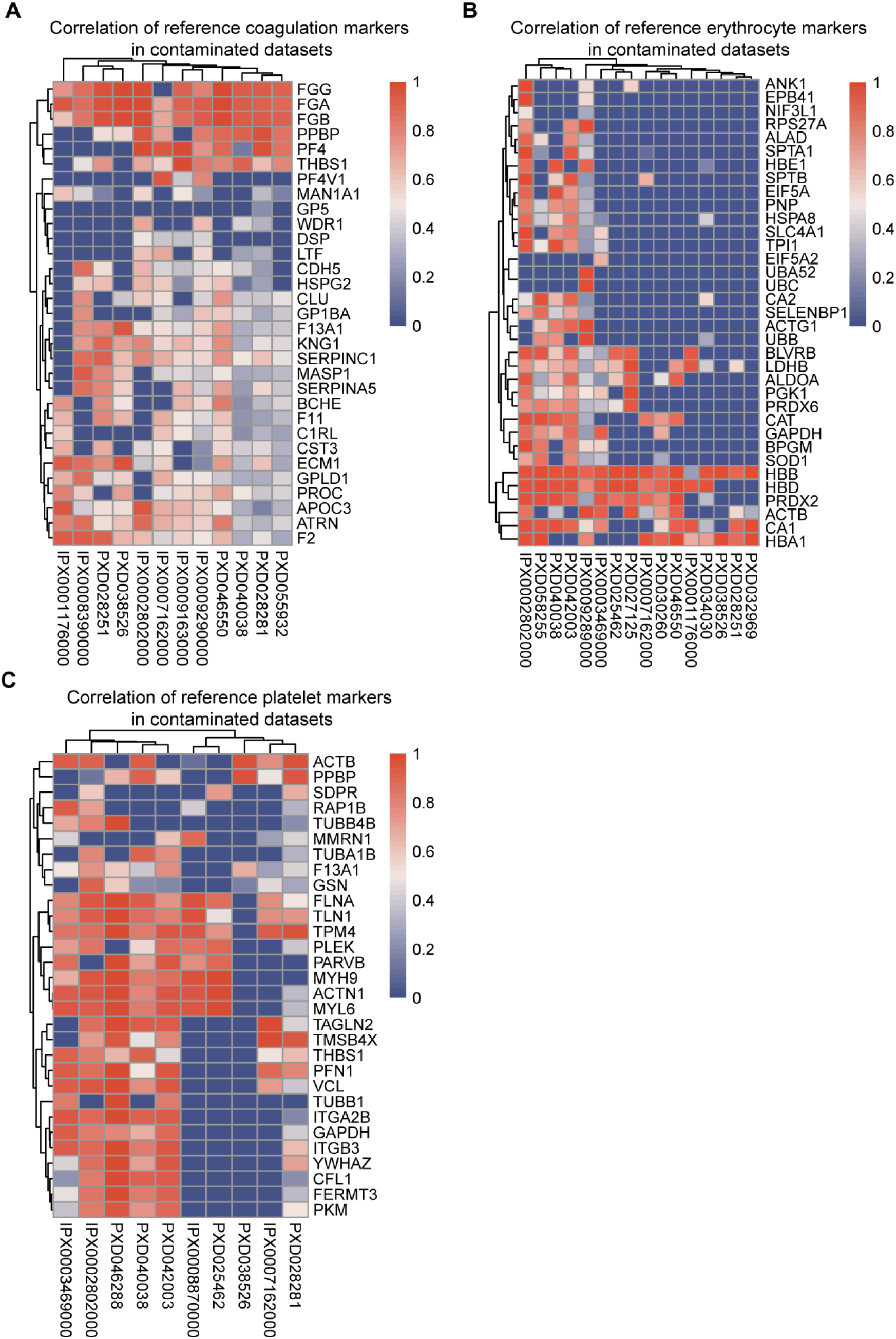
Evaluation of the suitability of experimentally validated contamination markers across distinct contaminated datasets. (A-C) Heatmaps showing the maximum correlation coefficients for platelet (A), erythrocyte (B), and coagulation (C) contamination markers across multiple contaminated datasets.

